# Evaluation of protein-ligand docking methods on peptide-ligand complexes for docking small ligands to peptides

**DOI:** 10.1101/212514

**Authors:** Sandeep Singh, Hemant Kumar Srivastava, Gaurav Kishor, Harinder Singh, Piyush Agrawal, G.P.S. Raghava

## Abstract

In the past, many benchmarking studies have been performed on protein-protein and protein-ligand docking however there is no study on peptide-ligand docking. In this study, we evaluated the performance of seven widely used docking methods (AutoDock, AutoDock Vina, DOCK 6, PLANTS, rDock, GEMDOCK and GOLD) on a dataset of 57 peptide-ligand complexes. Though these methods have been developed for docking ligands to proteins but we evaluate their ability to dock ligands to peptides. First, we compared TOP docking pose of these methods with original complex and achieved average RMSD from 4.74Å for AutoDock to 12.63Å for GEMDOCK. Next we evaluated BEST docking pose of these methods and achieved average RMSD from 3.82Å for AutoDock to 10.83Å for rDock. It has been observed that ranking of docking poses by these methods is not suitable for peptide-ligand docking as performance of their TOP pose is much inferior to their BEST pose. AutoDock clearly shows better performance compared to the other six docking methods based on their TOP docking poses. On the other hand, difference in performance of different docking methods (AutoDock, AutoDock Vina, PLANTS and DOCK 6) was marginal when evaluation was based on their BEST docking pose. Similar trend has been observed when performance is measured in terms of success rate at different cut-off values. In order to facilitate scientific community a web server PLDbench has been developed (<u>http://webs.iiitd.edu.in/raghava/pldbench/</u>).

## INTRODUCTION

Docking methods are widely used to study the molecular interactions between receptor and ligand molecules, thereby facilitating the process of drug discovery [1,2]. There are several well-established docking methods for the prediction of protein-protein [3–8], nucleic acid-ligand [9–11] and protein-ligand [12–17] interactions. In the past, many benchmarking studies have been carried out to evaluate the performance of docking methods and scoring functions [18–22]. However, benchmarking studies on docking methods involving peptide-ligand interactions are not available in the literature.

Peptide-ligand interactions have important applications in the treatment of diseases [23–26], development of drug-delivery systems [24,27], diagnostics [28,29] as well as in the development of sensor devices for rapid and reliable measurement of the concentration of target molecules [30]. In Alzheimer’s disease, small molecules have been utilized as potential inhibitors, which prevent the oligomerization and aggregation of amyloid β peptides [23]. Rodriguez et al designed and developed multifunctional thioflavin-based small molecules, which served as molecular probes in the detection of peptide amyloid fibrils as well as therapeutic agents in the treatment of Alzheimer’s disease [28]. Peptides such as cell-penetrating peptides (CPPs) are widely used as drug delivery vehicles to deliver small molecule drugs inside the cell [31–33]. Small molecules bind to CPPs by either covalent or non-covalent interaction and are transported into the cell [34]. Recently, the combination therapy utilizing the combination of cell-penetrating peptides and small molecule antibiotics has been proposed as a potential alternative in the treatment of infections caused by methicillin-resistant *Staphylococcus aureus* [24]. In recent years, peptide-based therapeutic is gaining attention due to their low toxicity and high specificity [35,36]. Therefore, many peptide-based resources have been developed in the last decade to enhance peptide-based therapeutics [37–46]. Thus, modeling peptide-ligand interactions require a detailed understanding of the way in which the small molecules bind to the active site of peptides.

Unfortunately, systematic parameterization in the docking methods is not available for studying peptide-ligand interactions. Docking methods efficiently produce acceptable docked poses but usually fail in ranking of the poses probably because of the failure of scoring function [47]. Development of a new scoring function is sometimes necessary for modeling the interactions of a new class of compounds. However, such lengthy and time-consuming process may not be essential if the available methods are capable of predicting the interactions of a new class of compounds. Best of author’s knowledge, no benchmarking study of the docking methods is currently available for modeling peptide-ligand interactions in the literature. We have benchmarked seven existing protein-ligand docking methods for their ability to correctly predict the peptide-ligand interactions on a series of 57 peptide-ligand complexes in this study. AutoDock [48], AutoDock Vina [16], DOCK 6 [49,13], PLANTS [14], rDock [10], GEMDOCK [15] and GOLD [17,50] docking methods were chosen for the benchmarking study. All of them have their strategies for the prediction of the best conformation of the ligand within the active site of a protein. All of these docking methods are available freely for academic use except GOLD. The extensive benchmarking of all the methods is carried out by analyzing 1, 3, 5, 10, 20 and 30 docked poses separately. We also tested the performance of tools like add hydrogen command of Open Babel [51] and Schrodinger [52].

## MATERIALS AND METHODS

### Benchmarking dataset of peptide-ligand complexes

All the peptide-ligand complexes were extracted from the Protein Data Bank (PDB) [53] based on the fulfillment of the following conditions: (i) length of peptide should be between 9 and 36 residues (9756 PDB entries), (ii) if the complex PDB is determined using X-RAY, it should have resolution < 2.5Å (2713 entries), (iii) ligand should not have any metal atoms (1748 entries), (iv) complexes having any covalent interactions between peptide and ligand atoms (as defined by distance between any peptide and ligand atom to be < 2Å) were removed using LPC software [54] (859 entries), (v) only one complex was selected if the peptide sequences of multiple complexes were identical (192 entries), (vi) complexes with only one ligand associated with its peptide were selected (75 entries). Finally, all the 75 entries were manually inspected and unusual entries (e.g. a single atom like Iodine being considered as ligand and entries with errors in docking calculations) were removed to get a dataset of 57 peptide-ligand complexes. The details of all the selected complexes are given in Table 1.

**Table 1:**
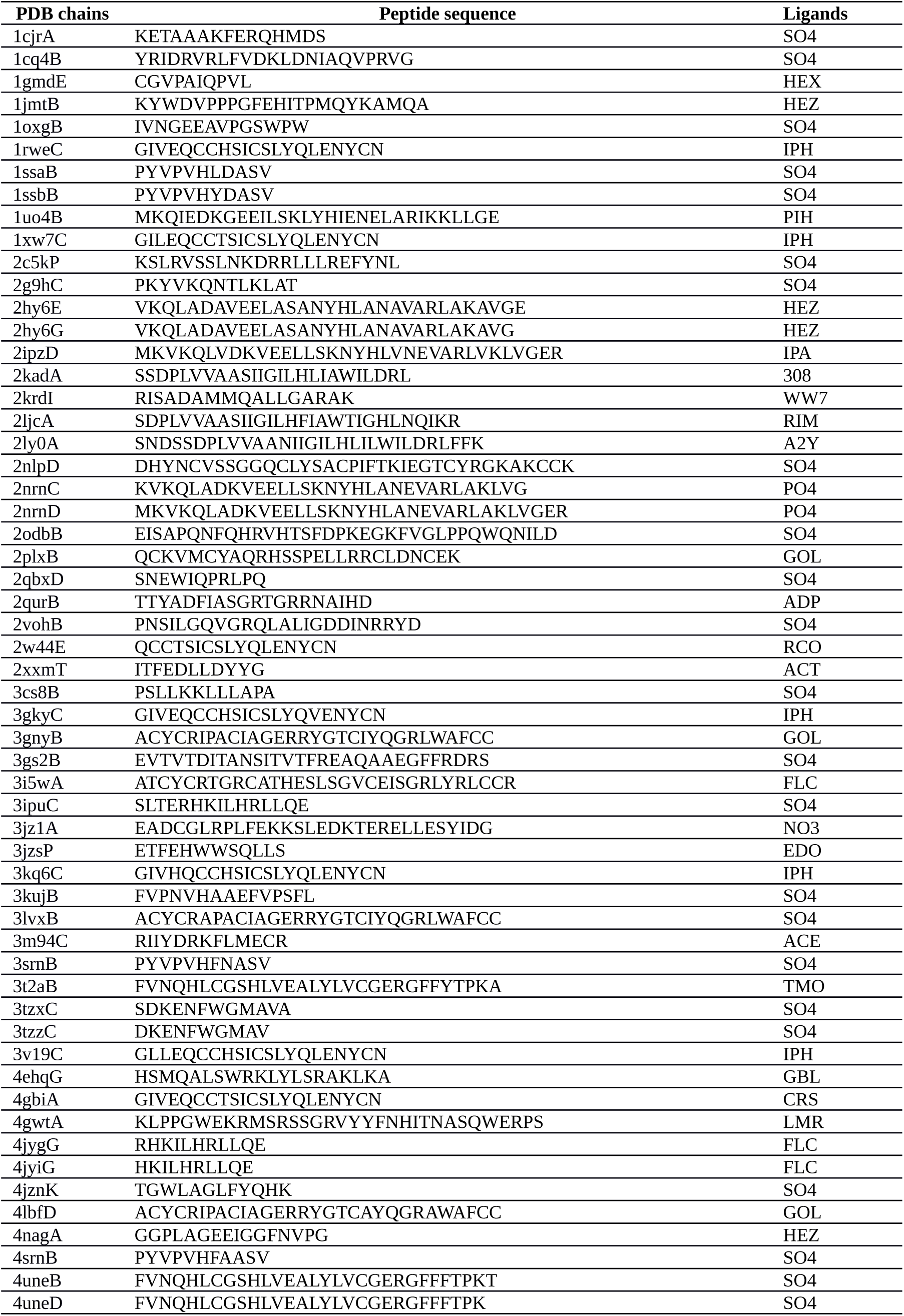
List of 57 peptide-ligand complexes used in this study for benchmarking, it includes amino acid sequence of peptides and name of ligand.

### Preprocessing of ligands and peptides

The ligands were extracted from the crystal structures and hydrogen atoms were added explicitly by using ‘add hydrogen tool’ available in Schrodinger module. The added hydrogen atoms were manually verified and errors (if any) were corrected. We used three initial geometries of ligands (first geometry is the coordinates of ligands as available in the PDB structures (represented as ‘crystal ligands’), second geometry is the energy minimized coordinates using GAFF force field in Open Babel (represented as ‘minimized ligands’) and third geometry is the ideal coordinates downloaded from Chemical Component Dictionary available in PDB (represented as ‘CCD ligands’)) for docking studies in order to see the effect of initial geometries of ligands on the overall results. The docking results of all the three geometries were analyzed separately. The peptides were extracted from their respective complexes and hydrogen atoms were added explicitly. All the water molecules were removed from the peptides before docking.

### Defining the binding site of the peptide-ligand interaction

Different docking methods define binding site either by creating a 3D-grid of X, Y and Z dimensions or by creating a sphere of a given radius centered at a defined point. In our study, we defined the center point by calculating the center of mass of the ligand molecule. We defined grid dimensions as a cube of length 40Å and radius of sphere 25Å. In this way, it was ensured that the search space in both the cases is approximately the same.

### Evaluation criteria

The root-mean-square deviation (RMSD) is a measure of the average distance between the atoms of superimposed structures. RMSD value is a widely used parameter to rank the performance of docking methods. If the docked ligand shows <2.0Å RMSD value with the crystallographic ligand, it is considered as a successful docking [18]. We calculated the RMSD values between docked pose and crystallographic pose using DOCK 6 [55]. DOCK 6 provides three types of RMSD values namely standard heavy atom RMSD, minimum-distance heavy atom RMSD and symmetry-corrected heavy atom RMSD. In our study, we used symmetry-corrected heavy atom RMSD, which is based on Hungarian algorithm. The Hungarian algorithm performs one-to-one assignments between original and docked ligand atoms and calculates the minimum distance between them [55].

### Docking methods

The ligands were docked to their respective peptides using seven different docking methods (AutoDock, AutoDock Vina, DOCK 6, PLANTS, rDock, GEMDOCK and GOLD). For each docking method, we used default parameters except defining the binding site as described in the above section. In order to evaluate the scoring function of the considered docking methods, we generated 30 docked poses on each docking method (except AutoDock Vina and GEMDOCK where maximum 20 poses were obtained). The results are analyzed and presented on the basis of 10, 20 and 30 poses separately. The docked pose with the top score (best score) is designated as ‘TOP pose’ and the docked pose having least RMSD value with the original pose is designated as ‘BEST pose’ in this study. A brief description of the docking methods used in this benchmark study is given below.

#### AutoDock

AutoDock is a frequently used and one of the most cited docking program in the scientific community. AutoDock uses Lamarckian genetic algorithm (LGA) for the prediction of the best conformation of a ligand within the active site of a receptor. It has empirical scoring function comprising van der Waals, electrostatic, hydrogen bonding and desolvation terms.

#### AutoDock Vina

AutoDock Vina is developed by the same group, who developed AutoDock. With a new scoring function, they improved the speed and accuracy of docking and compared it with AutoDock 4. AutoDock Vina is based on Iterated Local Search global optimizer algorithm for searching conformational space. Moreover, it is compatible with the PDBQT file format, which is used by AutoDock.

#### DOCK 6

DOCK 6 uses incremental construction approach and tries to geometrically match the ligand atoms within the receptor-binding site. It uses the energy-based scoring function as well as a grid-based footprint scoring function, which is used in rescoring the poses after docking.

#### PLANTS

PLANTS (Protein-Ligand ANT System), a new docking method, is based on ant colony optimization (ACO) algorithm. ACO mimics the behavior of real ants, which try to find the shortest path from their nest to a food source. The ants mark the path between the nest and food resource for their communication. In protein-ligand docking, an artificial ant colony is used to find minimum energy conformation of a ligand in the binding site.

#### rDock

rDock is originated from a program called RiboDock [11], developed for virtual screening of RNA targets. It uses stochastic as well as deterministic search techniques implementing genetic algorithm, Monte Carlo and simplex minimization stages for searching conformational space. The scoring function of rDock includes intermolecular terms like van der Waals forces and polar desolvation. rDock consist of two main programs *rbcavity* for the cavity generation and *rbdock* for the docking.

#### GEMDOCK

Generic Evolutionary Method for molecular DOCking (GEMDOCK) uses global as well as local search strategies to search for the conformational space of the ligand and uses empirical scoring function to score the generated poses. The scoring function of GEMDOCK uses energy parameters like steric, electrostatic and hydrogen bonding potentials.

#### GOLD

Genetic Optimization for Ligand Docking (GOLD) is one of the very promising molecular docking tools. It uses genetic algorithm for searching the conformational space of the ligand and docking it into the receptor-binding site. GOLD works on the basis of a fundamental requirement that the ligand molecule should have the capability to displace water molecules, which are loosely bound to the receptor.

## RESULTS AND DISCUSSION

This study is conducted to evaluate seven widely used protein-ligand docking methods (six of them are freely available for academic use except GOLD) for their ability to successfully model the peptide-ligand interactions. 57 different peptide-ligand complexes (fitted to our selection criteria as explained in the methodology section) were used as a dataset for this purpose.

### Performance of docking methods based on TOP docking poses

The performance of all docking methods computed on the basis of their TOP docking pose reveals that AutoDock performed the best while GEMDOCK performed worst with average RMSD 4.735Å and 12.627Å respectively (Table 2). AutoDock showed 21.05% success in reproducing crystallographic poses within 2Å RMSD and rest of the docking methods showed much less success rate (Table 2). As shown in Table 3, the success rate of docking methods improved as we increased cutoff value of RMSD. The performance of each docking pose of different methods on each peptide-ligand complex has been shown in supplementary tables (Table S1-S7). It was observed that most of the methods performed worst on certain peptide-ligand complexes. Therefore we assigned these peptide-ligand complexes as outliers and evaluated the performance of docking methods after removing outliers. The number of outliers at an RMSD cutoff of 8.5Å and 8Å were 2 and 6 respectively. The performance of docking methods on all complexes, after removing 2 outliers and after removing 6 outliers is shown in Table S8. The performance of most of the methods improved significantly after removing outliers. TOP docking pose obtained from any docking software may or may not be the BEST docking pose (having least RMSD with original pose). Therefore, we also compared the performance of these docking methods in terms of generating the BEST docking pose.

**Table 2:** The performance of different docking methods on 57 peptide-ligand complexes in terms of RMSD between docked and original poses as well as success rate. These methods were evaluated on the basis of their TOP and BEST docking pose.

| Docking methods | Performance of TOP and BEST docking pose |  |
| --- | --- | --- |
|  | RMSD | % Success |
| AutoDock-30 | 3.816 | 35.09 |
| AutoDock-20 | 3.956 | 31.58 |
| AutoDock-10 | 4.230 | 28.07 |
| AutoDock-5 | 4.505 | 26.32 |
| AutoDock-3 | 4.655 | 22.81 |
| AutoDock-1 | 4.735 | 21.05 |
| AutoDock Vina-20 | 4.496 | 40.35 |
| AutoDock Vina-10 | 6.601 | 28.07 |
| AutoDock Vina-5 | 8.806 | 14.04 |
| AutoDock Vina-3 | 9.528 | 10.53 |
| AutoDock Vina-1 | 10.420 | 10.53 |
| GEMDOCK-20 | 9.531 | 8.77 |
| GEMDOCK-10 | 10.477 | 8.77 |
| GEMDOCK-5 | 10.940 | 7.02 |
| GEMDOCK-3 | 11.795 | 7.02 |
| GEMDOCK-1 | 12.627 | 7.02 |
| rDock-30 | 10.826 | 5.26 |
| rDock-20 | 11.280 | 5.26 |
| rDock-10 | 11.558 | 5.26 |
| rDock-5 | 11.786 | 5.26 |
| rDock-3 | 11.869 | 5.26 |
| rDock-1 | 12.300 | 5.26 |
| PLANTS-30 | 4.480 | 43.86 |
| PLANTS-20 | 4.872 | 43.86 |
| PLANTS-10 | 6.682 | 35.09 |
| PLANTS-5 | 8.773 | 24.56 |
| PLANTS-3 | 9.595 | 21.05 |
| PLANTS-1 | 11.299 | 12.28 |
| GOLD-30 | 7.251 | 21.05 |
| GOLD-20 | 8.045 | 15.79 |
| GOLD-10 | 8.575 | 12.28 |
| GOLD-5 | 9.341 | 10.53 |
| GOLD-3 | 10.469 | 8.77 |
| GOLD-1 | 11.669 | 3.51 |
| DOCK 6-30 | 4.142 | 22.81 |
| DOCK 6-20 | 4.760 | 22.81 |
| DOCK 6-10 | 6.071 | 15.79 |
| DOCK 6-5 | 6.922 | 10.526 |
| DOCK 6-3 | 7.732 | 5.263 |
| DOCK 6-1 | 8.961 | 5.26 |

RMSD is the root mean square deviation between docked and crystallographic poses. % Success is the percent success rate in reproducing the docked poses within 2.0Å RMSD with the original pose. 3, 5, 10, 20 and 30 notations show the number of generated poses to select the BEST pose. 1 notation shows the TOP pose.

**Table 3:** The performance of different docking methods in terms of percent success rate at different RMSD cut-off.

| **RMS**<br>**D**<br>**(Å)** | **% Success rate of TOP docking pose** |  |  |  |  |  |  |
| --- | --- | --- | --- | --- | --- | --- | --- |
|  | **AutoDock**<br>**k** | **AutoDock Vina** | **GEMDOCK** | **rDock** | **PLANTS** | **GOLD** | **DOCK 6** |
| 2.00 | 21.05 | 10.53 | 7.02 | 5.26 | 12.28 | 3.51 | 5.26 |
| 2.25 | 31.58 | 10.53 | 7.02 | 5.26 | 15.79 | 7.02 | 7.02 |
| 2.50 | 35.09 | 10.53 | 8.77 | 5.26 | 19.30 | 8.77 | 7.02 |
| 2.75 | 36.84 | 10.53 | 10.53 | 7.02 | 19.30 | 10.53 | 8.77 |
| 3.00 | 40.35 | 10.53 | 10.53 | 7.02 | 21.05 | 12.28 | 8.77 |
| 4.00 | 45.61 | 12.28 | 12.28 | 7.02 | 24.56 | 15.79 | 8.77 |
| 5.00 | 50.88 | 14.04 | 12.28 | 10.53 | 26.32 | 15.79 | 12.28 |
| 8.00 | 84.21 | 28.07 | 21.05 | 17.54 | 33.33 | 21.05 | 35.09 |
| 10.00 | 98.25 | 57.89 | 38.60 | 33.33 | 42.11 | 35.09 | 59.65 |
| 12.00 | 100 | 73.68 | 57.89 | 52.63 | 56.14 | 52.63 | 84.21 |
| 15.00 |  | 82.46 | 71.93 | 68.42 | 66.67 | 71.93 | 98.25 |
| 20.00 |  | 87.72 | 80.70 | 89.47 | 85.96 | 94.74 | 98.25 |
| 25.00 |  | 100 | 92.98 | 100 | 100 | 98.25 | 100 |
| 30.00 |  |  | 98.25 |  |  | 100 |  |
| 35.00 |  |  | 100 |  |  |  |  |
| **RMS**<br>**D**<br>**(Å)** | **% Success rate of BEST docking pose** |  |  |  |  |  |  |
|  | **AutoDock**<br>**k** | **AutoDock Vina** | **GEMDOCK** | **rDock** | **PLANTS** | **GOLD** | **DOCK 6** |
| 2.00 | 35.09 | 40.35 | 8.77 | 5.26 | 43.86 | 21.05 | 22.81 |
| 2.25 | 47.37 | 43.86 | 10.53 | 8.77 | 43.86 | 24.56 | 33.33 |
| 2.50 | 47.37 | 43.86 | 12.28 | 10.53 | 47.37 | 24.56 | 38.60 |
| 2.75 | 47.37 | 47.37 | 12.28 | 10.53 | 50.88 | 26.32 | 42.11 |
| 3.00 | 50.88 | 47.37 | 14.04 | 10.53 | 54.39 | 26.32 | 43.86 |
| 4.00 | 59.65 | 52.63 | 15.79 | 12.28 | 57.89 | 31.58 | 57.89 |
| 5.00 | 61.40 | 61.40 | 21.05 | 12.28 | 57.89 | 36.84 | 64.91 |
| 8.00 | 89.47 | 80.70 | 38.60 | 26.32 | 82.46 | 56.14 | 94.74 |
| 10.00 | 100 | 89.47 | 63.16 | 45.61 | 91.23 | 73.68 | 96.49 |
| 12.00 |  | 98.25 | 77.19 | 66.67 | 96.49 | 80.70 | 98.25 |
| 15.00 |  | 100 | 87.72 | 78.95 | 98.25 | 96.49 | 100 |
| 20.00 |  |  | 92.98 | 92.98 | 100 | 100 |  |
| 25.00 |  |  | 96.49 | 100 |  |  |  |
| 30.00 |  |  | 98.25 |  |  |  |  |
| 35.00 |  |  | 100 |  |  |  |  |

TOP pose means the pose with best docking score. BEST pose means the pose having lowest RMSD value compared to the original pose.

### Performance of docking methods based on BEST docking poses

In addition to TOP docking pose, we identified and evaluated BEST docking pose from number of docking poses generated by different methods. AutoDock Vina and GEMDOCK generated maximum 20 docking poses wereas for the rest of the docking methods, upto 30 docking poses were generated corresponding to each ligand. The performance of BEST docking pose among top 3, 5, 10, 20 and 30 poses generated by different methods is shown in Table 2. AutoDock showed 22.81%, 26.32%, 28.07%, 31.58% and 35.09% success rate for the top 3, 5, 10, 20 and 30 poses respectively. The average RMSD also decreased (4.655Å for 3 poses, 4.505Å for 5 poses, 4.230Å for 10 poses and 3.816Å for 30 poses) with the increase of the docked poses in AutoDock. AutoDock Vina and PLANTS docking methods showed the most dramatic effect on the overall success rate with increase in the number of poses. Success rate increased from 28.07% to 40.35% in AutoDock Vina and 35.09% to 43.86% in PLANTS as the docked poses increased from 10 to 20. Success rate for PLANTS was close to the success rate for AutoDock on the basis of 3 and 5 docked poses. Noticiable improvement in the success rate was obtained in AutoDock Vina and PLANTS on the increase of the number of docked poses from 1 to 10 (10.53% to 28.07% for AutoDock Vina and 12.28% to 35.09% for PLANTS). AutoDock Vina and PLANTS showed more than 40% success rate for 20 poses which is better than the success rate of Audodock for the same number of poses. This clearly indicates that AutoDock Vina and PLANTS docking methods are able to generate many docked poses close to the crystallographic pose, however their scoring function is not able to give them higher scores. Average RMSD values reduced from 6.601Å to 4.496Å for AutoDock Vina and 6.682Å to 4.872Å for PLANTS while increasing the docked poses from 10 to 20. Most of the docking methods showed improved RMSD values even for 3 and 5 docked poses. However, the improvement was highest in PLANTS (11.299Å for 1 pose and 8.773Å for 5 poses) and lowest in AutoDock (4.735Å for 1 pose and 4.505Å for 5 poses). Much improvement in the RMSD values were obtained on the increase of the number of poses from 1 to 10 in AutoDock Vina and PLANTS. Success rate and average RMSD values were not much affected with the increase in the docked poses in rDock and GEMDOCK docking methods. Therefore, rDock and GEMDOCK methods completely fail to model peptide-ligand interactions. GOLD and DOCK 6 docking methods showed some improvement in the success rate with the increase in the docked poses but the results are not better than AutoDock, AutoDock Vina or PLANTS. DOCK 6 showed reasonable average RMSD values while considering 30 docked poses. The lowest RMSD values for each ligand for all the docking methods are depicted in Table S9; clearly AutoDock shows lower RMSD values for most of the ligands as compared to other docking methods. Figure 1 depicts the RMSD variation in the TOP and BEST poses for all the docking methods. The RMSDs of all the docking methods are arranged in the increasing order to clearly see the variation and thus they are irrespective of the PDB-IDs. The RMSD variations of the best-of-TOP poses and best-of-BEST poses are also included in Figure 1. Best-of TOP was obtained by taking the best values provided by any docking method for the TOP poses and best-of-BEST was obtained by taking the best values provided by any docking method for the BEST poses. Clearly, the RMSD values obtained from best-of-TOP and the RMSD values obtained from AutoDock is almost similar. However, AutoDock, AutoDock Vina, PLANTS and DOCK 6 docking methods show similar trends for the BEST poses. The best-of-BEST shows much improved performance and indicates that the combination of existing methods may provide reasonable success rate for modeling peptide-ligand interactions. Figure 2 depicts the RMSD variation in the top 20 poses for all the docking methods. The RMSD values of >5.0Å are not included in Figure 2 for better representation of the data. All the values are provided in the Table S1-S7 for details. Figure 2 clearly indicates that AutoDock generates most of the poses in lower RMSD range and same is not true for other docking methods. Figure 3 shows a case study (1xw7C), depicting the TOP and BEST docked poses obtained from various docking methods.

**Figure 1:**
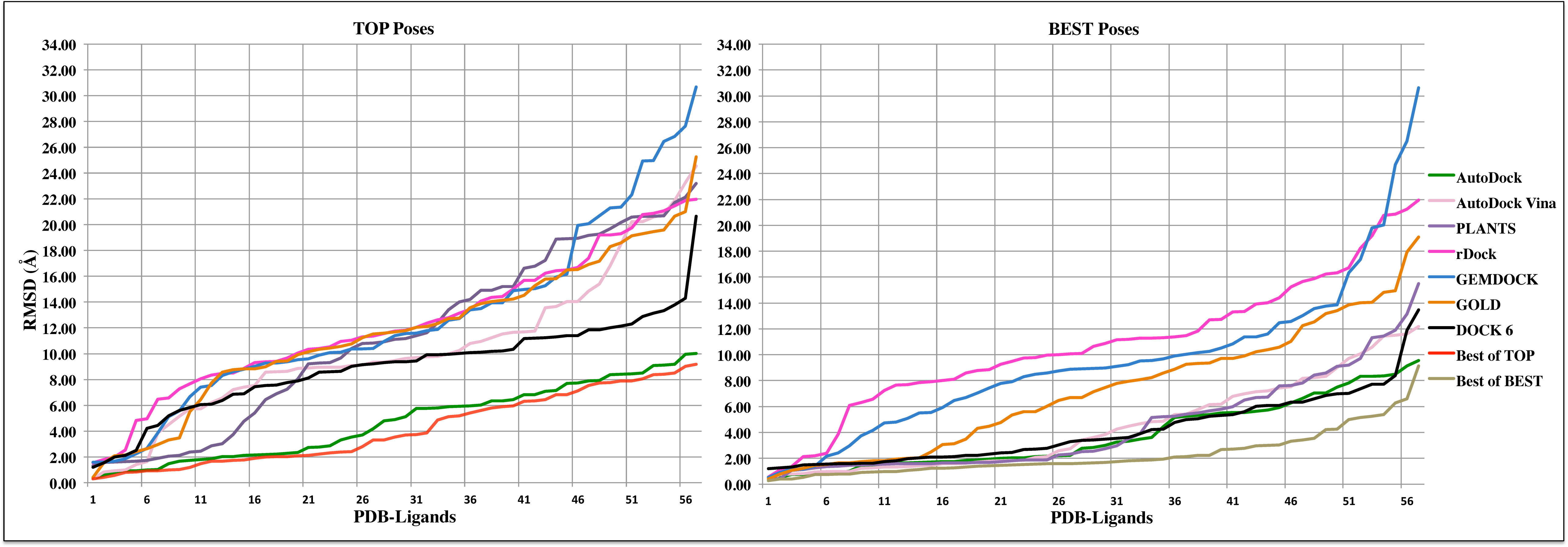
The performance (RMSD between docking and original pose) of different methods based on their TOP and BEST docking pose, including best-of-TOP poses and best-of-BEST poses.

**Figure 2:**
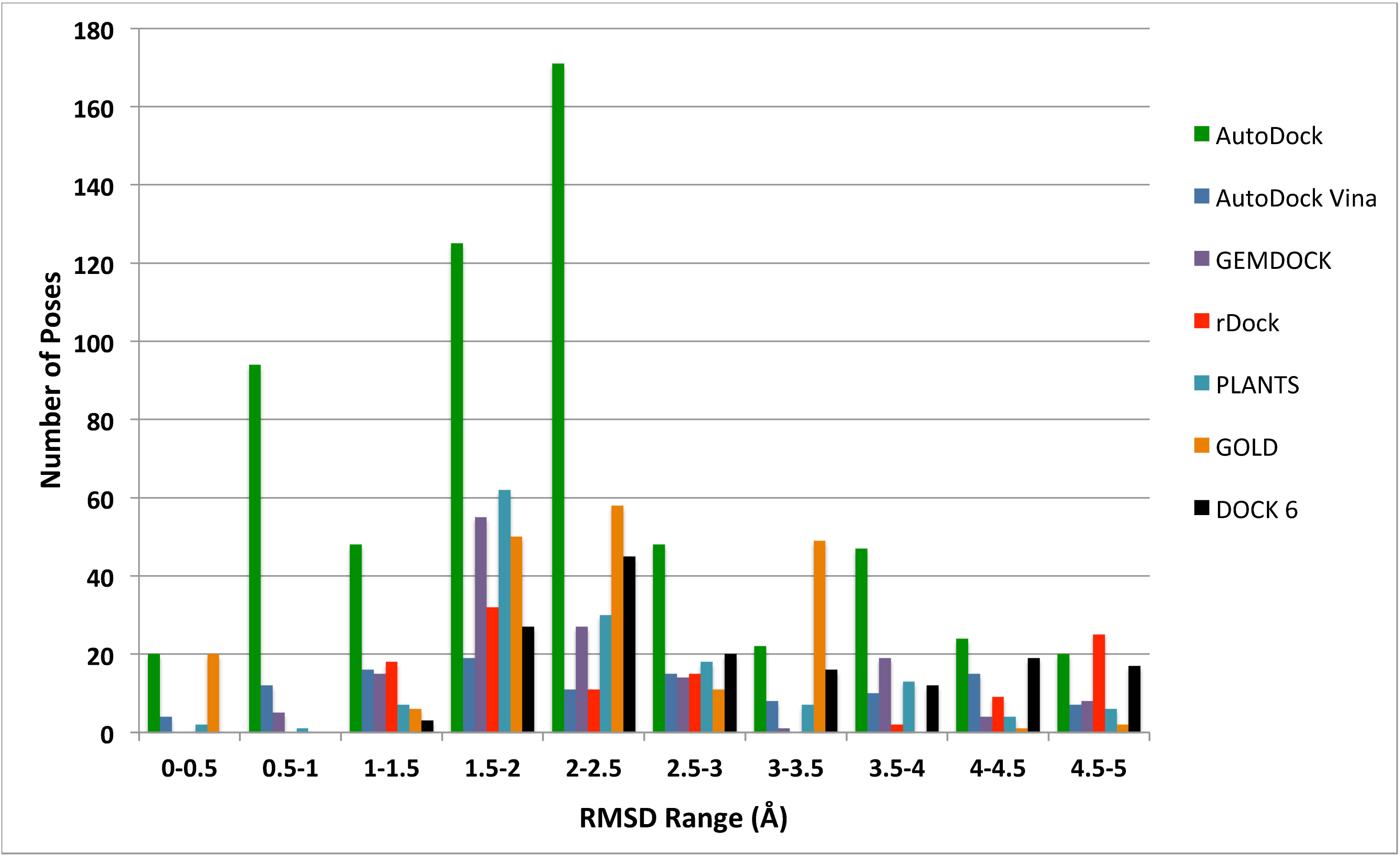
RMSD variation in top 20 poses for all the docking methods.

**Figure 3:**
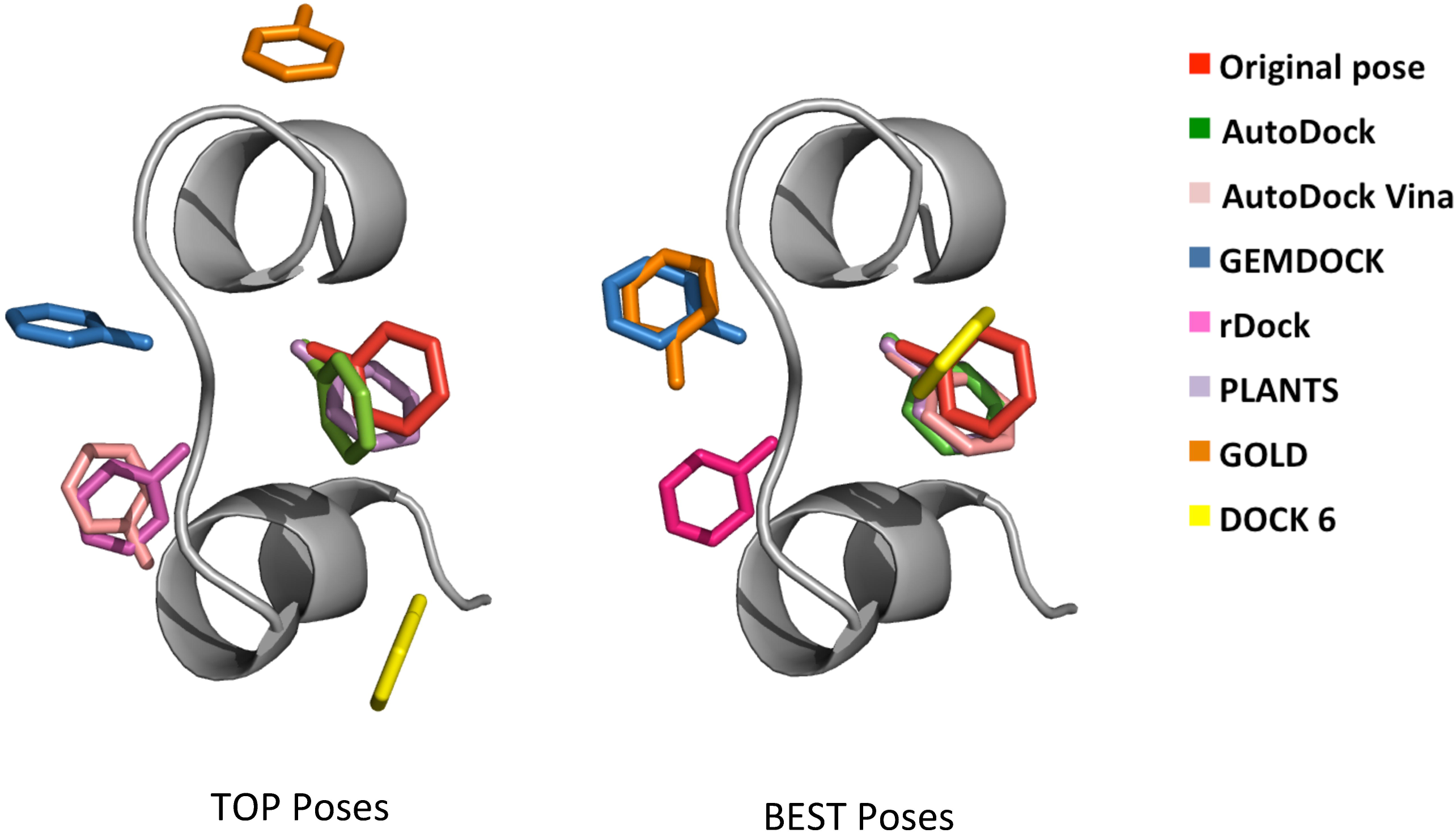
A case study (1xw7C) of the comparison of docked poses obtained from various docking methods.

Table 3 shows the success rate on the basis of sequential increase in cutoff RMSD values. Interestingly the success rate of some of these methods increases dramatically in producing the docked poses a little higher than 2.0Å RMSD with the original pose. AutoDock, AutoDock Vina, PLANTS, DOCK 6 and GOLD docking methods respectively show 47.37%, 43.86%, 43.86%, 33.33% and 24.56% success in producing the docked poses within 2.25Å RMSD with the crystallographic poses. The success rate of AutoDock, AutoDock Vina and PLANTS docking methods are comparable for 3.00Å cutoff value. GEMDOCK and rDock fails to produce acceptable success rate even after increasing the cutoff value from 2.0Å to 3.0Å. We also calculated the maximum possible success rate by combining the results obtained from all the docking methods. For this, we calculated success rate of best-of-TOP pose (lowest RMSD value achieved by the TOP poses of all the docking methods for each complex) and best-of-BEST pose (lowest RMSD value achieved by the BEST poses of all the docking methods for each complex). 61.40% and 75.40% success rate was achieved by best-of-TOP pose at 2.00Å and 3.00Å RMSD cutoff values respectively. In addition, much improvement was observed in the average RMSD value (2.30Å) for best-of-BEST pose. Clearly, if one can combine the scoring function of all these docking methods, ~75% success rate may be achieved and reasonable average RMSD values may be obtained. This directs the path of the future work in this area. Table 4 depicts the RMSD variation of TOP and BEST docking poses generated by all the docking methods. All the docking methods shows higher RMSD variations and hence are capable of generating diverse poses. RMSD variation while considering the BEST pose, is very high for GEMDOCK and rDock and is lowest for AutoDock.

**Table 4:** Variation in performance (average RMSD and RMSD range) of TOP and BEST docking poses generated by different docking methods on 57 peptide-ligand complexes.

| Docking methods | Average RMSD (RMSD Range) (Å) |  |
| --- | --- | --- |
|  | TOP poses | BEST poses |
| AutoDock | 4.735 (0.32 – 10.02) | 3.816 (0.28 – 9.56) |
| AutoDock Vina | 10.420 (0.30 – 24.55) | 4.496 (0.28 – 12.19) |
| GEMDOCK | 12.627 (1.59 – 30.68) | 9.531 (0.52 – 30.62) |
| rDock | 12.300 (1.53 – 21.97) | 10.826 (0.56 – 21.95) |
| PLANTS | 11.299 (1.31 – 23.19) | 4.480 (0.40 – 15.50) |
| GOLD | 11.669 (0.41 – 25.25) | 7.251 (0.41 – 19.08) |
| DOCK 6 | 8.961 (1.20 – 20.64) | 4.142 (1.20 – 13.47) |

TOP pose means the pose with best docking score. BEST pose means the pose having lowest RMSD value compared to the original pose.

### Evaluation of scoring capability of docking methods

The foremost purpose of any docking method is to differentiate between the true solutions (usually defined as the ones docked within 2.0Å RMSD from the original structure) and misdocked structures. This differentiation is based on the scoring function of the docking methods. Thus, scoring function is very crucial to get correct docking results. We benchmarked the scoring function of all the docking methods on our dataset by generating various docked poses. Table 5 shows the performance of scoring function and extent of deviation in the scoring of all the docking methods. Any successful docking should either produce the TOP pose as the BEST pose or should show the minimum deviation among them. All the docking methods show poor success in producing the same pose as TOP pose and BEST pose. However, AutoDock shows 91.23% success in producing the TOP and the BEST poses within the difference of 0.75Å RMSD while considering 10 generated docked poses. Success rate in producing the TOP pose and the BEST pose within minimum deviation is higher for rDock, GEMDOCK and GOLD, and lower for PLANTS, AutoDock Vina and DOCK 6. Moreover, the significant difference in average RMSD values between TOP pose and BEST pose in AutoDock Vina and PLANTS shows that their scoring function are unable to give high scores to BEST pose (Table 2). Overall, the scoring function of AutoDock is able to model the peptide-ligand complexes better than other methods on our dataset.

**Table 5:** Percent of cases with absolute difference between BEST and TOP poses at different RMSD cut-off.

| Docking methods | RMSD cut-offs |  |  |  |
| --- | --- | --- | --- | --- |
| | $\vert B-T\vert =0\text{\AA}$ | $\vert B-T\vert \leq 0.25\text{\AA}$ | $\vert B-T\vert \leq 0.50\text{\AA}$ | $\vert B-T\vert \leq 0.75\text{\AA}$ |
| AutoDock-30 | 1.75 | 52.63 | 70.18 | 77.19 |
| AutoDock-20 | 1.75 | 59.65 | 75.44 | 84.21 |
| AutoDock-10 | 5.26 | 73.68 | 84.21 | 91.23 |
| AutoDock-5 | 19.30 | 84.21 | 91.23 | 96.49 |
| AutoDock-3 | 35.09 | 89.47 | 94.74 | 98.25 |
| AutoDock Vina-20 | 5.26 | 15.79 | 17.54 | 19.30 |
| AutoDock Vina-10 | 10.53 | 29.82 | 35.09 | 38.60 |
| AutoDock Vina-5 | 22.81 | 56.14 | 63.16 | 64.91 |
| AutoDock Vina-3 | 50.88 | 78.95 | 80.70 | 84.21 |
| GEMDOCK-20 | 1.75 | 40.35 | 50.88 | 56.14 |
| GEMDOCK-10 | 1.75 | 52.63 | 57.89 | 63.16 |
| GEMDOCK-5 | 15.79 | 63.16 | 68.42 | 68.42 |
| GEMDOCK-3 | 31.58 | 75.44 | 77.19 | 78.95 |
| rDock-30 | 3.51 | 43.86 | 54.39 | 63.16 |
| rDock-20 | 8.77 | 57.89 | 64.91 | 71.93 |
| rDock-10 | 12.28 | 63.16 | 71.93 | 82.46 |
| rDock-5 | 19.30 | 78.95 | 80.70 | 87.72 |
| rDock-3 | 33.33 | 85.96 | 89.47 | 92.98 |
| PLANTS-30 | 3.51 | 10.53 | 12.28 | 15.79 |
| PLANTS-20 | 3.51 | 10.53 | 14.04 | 17.54 |
| PLANTS-10 | 10.53 | 21.05 | 29.82 | 35.09 |
| PLANTS-5 | 21.05 | 38.60 | 50.88 | 61.40 |
| PLANTS-3 | 38.60 | 57.89 | 66.67 | 70.18 |
| GOLD-30 | 3.51 | 14.04 | 21.05 | 24.56 |
| GOLD-20 | 8.77 | 26.32 | 33.33 | 36.84 |
| GOLD-10 | 14.04 | 31.58 | 38.60 | 42.11 |
| GOLD-5 | 22.81 | 42.11 | 49.12 | 56.14 |
| GOLD-3 | 40.35 | 61.40 | 71.93 | 78.95 |
| DOCK 6-30 | 5.26 | 10.53 | 12.28 | 12.28 |
| DOCK 6-20 | 8.77 | 15.79 | 17.54 | 17.54 |
| DOCK 6-10 | 14.04 | 26.32 | 26.32 | 33.33 |
| DOCK 6-5 | 17.54 | 40.35 | 47.37 | 52.63 |
| DOCK 6-3 | 29.82 | 56.14 | 61.40 | 70.18 |

$|B-T|$ represents the absolute difference in the RMSD between the BEST and the TOP pose. 3, 5, 10, 20 and 30 notations show the number of generated poses to select the BEST pose.

### Effect of initial geometry of ligand on docking results

We prepared three different initial geometries of ligands for docking. First geometry is the coordinates of ligands as available in the PDB (crystal ligands), second geometry is obtained from energy minimization of PDB ligands using GAFF force field with the help of Open Babel program (minimized ligands) and third geometry is the coordinates of ligands downloaded from the Chemical Component Dictionary available in PDB (CCD ligands). Summary of docking results using all three initial geometries of ligands are given in Table 2, S10 and S11 respectively for crystal, minimized and CCD ligands. It is clear that the initial geometry of ligands is not affecting the overall success rate and average RMSD values. This may be associated with the size of the ligands as most of the ligands considered in this study are small in size.

### Correct addition of hydrogen atoms

Correctly adding hydrogen atoms in ligands and receptors is crucial in order to get the meaningful docking results. We tested the ‘add hydrogen’ tool of Open Babel and Schrodinger packages in adding the hydrogen atoms explicitly to the ligand molecules in our dataset. Schrodinger package shows 61% success while Open Babel shows 40% success in correctly adding hydrogen atoms in the considered ligands. The success rate of ‘add hydrogen’ tool of both the programs is given in Table S12. The performance of Schrodinger is better as compared to Open Babel in adding hydrogen atoms to the current dataset. However, both these tools are not able to correctly add hydrogen atoms on all the ligands. Generally, Open Babel fails to correctly add hydrogen atoms in charged ligands like SO_4_^2–^, PO_4_ ^3–^ etc. and Schrodinger generally fails in aromatic ligands like C_6_H_5_OH (phenol), C_6_H_6_O_2_ (resorcinol) etc. The failure of ‘add hydrogen tool’ of Schrodinger may be associated with the higher bond lengths in PDB structures. Thus, manual verification of added hydrogen in ligand molecules is necessary before proceeding to the next step in docking.

### Effect of charged ligands on overall docking results

We divided the whole dataset in charged and uncharged ligands in order to understand the effect of charge on ligand molecules on overall docking results. Table S13 shows the performance of all the considered docking methods for charged and uncharged ligands separately. Considering the TOP pose, AutoDock docking method shows 25% success rate for charged ligands and only 15.38% success rate for uncharged ligands. In fact the performance for all the considered docking methods, except DOCK-6, is better for charged ligands compared to uncharged ligands for both BEST and TOP docking poses. On the other hand DOCK-6 shows comparable performance for charged and uncharged ligands. The performance of AutoDock is much better compared to all other docking methods for TOP docking poses. Considering BEST docking poses, PLANTS shows the best performance with 53.12% success rate for charged and 34.62% for uncharged ligands. Interestingly, AutoDock Vina performs better for charged and uncharged ligands compared to AutoDock. In general, docking methods show better success rate for charged ligands.

### Effect of aromatic ligands on overall docking results

We tested the effect of aromatic ligands on overall docking results. The Table S14 depicts the docking results for aromatic and non-aromatic ligands for all the considered docking methods for both TOP and BEST poses. All the docking methods, except AutoDock, completely fail for aromatic ligands on the basis of TOP docking poses. AutoDock shows 27.27% success rate for aromatic and 19.15% success rate for other ligands for TOP poses and this is the best performance. AutoDock vina shows most dramatic effect as the success rate increases from 0% to 54.55% on considering BEST poses. Success rate is 0% for GOLD, rDOCK and GEMDOCK docking methods even for BEST docking poses on considering aromatic ligands. On the other hand, PLANTS, AutoDock Vina, AutoDock and DOCK 6 docking methods show better performance for aromatic ligands on considering BEST poses. Thus, AutoDock is much better compared to all other docking methods for aromatic ligands on the basis of TOP poses. Interestingly, the performance of AutoDock Vina docking method is best for aromatic ligands and PLANTS docking method for non aromatic ligands on the basis of BEST poses.

### Implementation of web server

Based on the above benchmarking study and to help the scientific community, we have developed a web server ‘PLDbench’ with an easy interface. This server has following modules: (i) Single: In this module a user can submit a docked pose and can select the original crystal ligand out of 57 peptide-ligand complexes (used in this benchmark study) from dropdown menu. User will get the RMSD value between these ligands. (ii) Benchmark: This module provides an option to perform a benchmark analysis on a set of docked poses given by a user in archived zip file format. (iii) Compare: This module provides an option to calculate RMSD between original ligand and predicted pose given by the user. PLDbench web service is freely accessible at <u>http://webs.iiitd.edu.in/raghava/pldbench/</u>

## LIMITATIONS

Currently no docking method is available specifically for modeling peptide-ligand interactions. Studying peptide-ligand interactions is difficult and suffers from following limitations: i). Peptides are more flexible than proteins and tend to adopt more than one conformation ii) binding of small molecules to peptides may trigger substantial conformational changes and therefore changing the tertiary structure of the peptide. iii)

Peptides lack a well-defined active site cavity and are generally involved in surface mediated interactions. Moreover, the peptide-ligand complexes extracted from the PDB and used as dataset in this study, lacks the structural diversity in small ligands, which further poses a limitation in studying peptide-ligand interactions. However, considering the importance of peptide-ligand docking, we need to understand this area even after these limitations. Thus, selection of methods from currently available docking methods may be crucial till a novel method is developed specifically for studying peptide-ligand interactions.

## CONCLUSION

The capability of 7 docking methods is benchmarked for their ability to model peptide-ligand interactions. Three different initial geometries of ligands were tested for docking. We observed that the docking results are not affected on the basis of initial geometries of ligand molecules. This work also indicates the necessity of manual verification of added hydrogen atoms in ligand molecules before proceeding to the next step in docking. AutoDock clearly shows much better performance compared to the other six docking methods to model peptide-ligand interactions considering the TOP pose. On the other hand, performance of AutoDock, AutoDock Vina, PLANTS and DOCK 6 are similar for the BEST poses. A slight increase in the cutoff values (>2.00Å to >2.25Å) shows much improvement in the success rate of most of the docking methods. Comparison of 3, 5, 10, 20 and 30 docked poses shows that 20 docked poses may be appropriate for modeling peptide-ligand interactions. Much improvement in the success rate and average RMSD values is observed on the combination of all the considered docking methods. Clearly, if one can combine the scoring function of all these docking methods, 75% success rate may be achieved and reasonable average RMSD values may be obtained for peptide-ligand interaction. This directs the path of the future work in this area. This work also shows the need to parameterize the docking methods for modeling peptide-ligand interactions and development of a new scoring function. However, one may use AutoDock for docking studies of peptide-ligand complexes till further advancements are achieved in this area.

## SUPPLEMENTARY MATERIAL

One file named ‘Supplementary file’ in PDF file format contains detailed information about RMSD values of all the docked poses generated by all the benchmarked docking methods.

## ACKNOWLEDGEMENTS

Authors’ are thankful to Council of Scientific and Industrial Research (CSIR), Department of Biotechnology (DBT) and Department of Science and Technology (DST), Government of India, for fellowships. GENESIS, BSC0121 and BTISNET are acknowledged for the financial support.

## Authors’ Contribution

HS collected and compiled the dataset. SS HKS GK performed the experiments. GK PA SS developed the web interface. HKS SS GK GPSR analyzed the data and prepared the manuscript. GPSR conceived the idea and coordinated the project.

## Conflict of Interest

The authors’ declare that they have no conflict of interest.

